# The transcription factor Pou3f1 provides a new map to the glutamatergic neurons of the cerebellar nuclei

**DOI:** 10.1101/2020.05.18.102855

**Authors:** Joshua Po Han Wu, Joanna Yeung, Sih-Rong Wu, Huda Zoghbi, Dan Goldowitz

## Abstract

Pou3f1 is a transcription factor involved in early neural differentiation. Cap Analysis Gene Expression (5’-CAGE) analysis reveals that *Pou3f1* transcript is highly enriched in the developing cerebellum. Between embryonic (E) days E10.5 and E12.5, Pou3f1 expression is present prominently along the subpial stream (SS), suggesting that Pou3f1^+^ cells are glutamatergic cerebellar nuclear (CN) neurons. This finding was confirmed by immunofluorescent (IF) co-labeling of Pou3f1 and Atoh1, the master regulator of cells from the rhombic lip (RL) that are destined for neurons of the glutamatergic lineage, as well as in *Atoh1*-null tissues, in which Pou3f1 expression is absent. Interestingly, the expression of Pax6, another key molecule for CN neuron survival, does not co-localize with that of Pou3f1. In the *Pax6-*null Small Eye (*Sey*) mutant, which is characterized by a loss of many glutamatergic CN neurons, Pou3f1^+^ CN neurons are still present. Furthermore, Pou3f1-labeled cells do not co-express Tbr1, a well-established marker of glutamatergic CN neurons. These results highlight that Pou3f1^+^ cells are a distinct and previously unrecognized subtype of glutamatergic CN neurons that do not have the “canonical” sequence of Atoh1→Pax6→Tbr1 expressions. Instead, they express Atoh1, Pou3f1, and other markers of CN neurons, Brn2 and Irx3. These findings illustrate that glutamatergic CN neurons that arise from the RL are composed of molecularly heterogeneous subpopulations that are determined by at least two distinct transcriptional programs.

**Significance Statement:** The present work has identified Pou3f1 as a marker for a previously unidentified subtype of glutamatergic cerebellar nuclear neurons, the principal output neurons of the cerebellum. The classical model of glutamatergic CN neurons development follows the sequential expression of transcription factors Atoh1→Pax6→Tbr1. However, we found that the development of Pou3f1^+^ neurons requires Atoh1 but not Pax6. Moreover, Pou3f1^+^ neurons do not express Tbr1, but instead express two other transcription factors, Brn2 and Irx3. Anatomically, Pou3f1^+^ CN neurons populate the interposed and dentate nuclei, while the Tbr1^+^ CN neurons populate the fastigial nucleus. These findings reveal the heterogeneity of CN neuron populations and the diversity of molecular programming that lead to different CN neuron subtypes.

## Introduction

The cerebellum is an ideal brain region for the study of neuronal development, owing to the limited number of cell types that construct the cerebellum and the well-defined developmental timeline for each cell type. Using this prior knowledge, we have been using the cerebellum as a proxy to gain insight into the molecular determinants of brain development. To elucidate the underlying gene regulatory cascades that produce cerebellar cell types, our lab in collaboration with the FANTOM5 Consortium collected a dynamic transcriptomic dataset on the cerebellum across embryonic and neonatal (P) timepoints (E11-P9) (Arner et al., 2015; Ha et al., 2019). Gene regulatory network analysis of this dataset has identified novel gene candidates that have not previously been associated with cerebellar development. One of these genes, *Pou3f1*, encodes a member of a family of transcription factors characterized by a POU DNA-binding domain, which consists of two highly conserved regions tethered by a variable linker (Ryan et al., 1997). Pou3f1 is highly expressed during gastrulation and in developing mouse brain. In addition, it has been suggested to play a role in neural differentiation (Dominguez et al., 2012). As demonstrated by a knockdown study in embryonic stem cells (ESCs) (Zhu et al., 2014), Pou3f1 acts primarily as a repressor of the BMP and Wnt signaling pathways. Expression levels of targets of the BMP pathway (Id1, Id2, Msx1, Msx2), and those of targets of the Wnt pathway (Wnt3, Axin2, Dkk1, and Myc), are significantly increased when *Pou3f1* is knocked down. Since both the BMP and Wnt signaling pathways are implicated in the maintenance of multipotency, by acting as a repressor, Pou3f1 likely plays a role in cell fate commitment. While there are accumulating data of the involvement of Pou3f1 in cortical development, such information is missing in the cerebellum.

Prior genetic fate mapping studies done on the cerebellum have yielded a spatial and genetic map in which two major cell lineages, the gamma-aminobutyric acidergic (GABAergic) and glutamatergic neurons, arise from two anatomically and molecularly distinct regions. The ventricular zone (VZ) that sits above the fourth ventricle is defined by the transcription factor Ptf1a, which gives rise to GABAergic populations, such as Purkinje cells, GABAergic interneurons, and GABAergic CN neurons (Hoshino et al., 2005). On the other hand, the RL is defined by the transcription factor Atoh1, which gives rise to glutamatergic populations such as granule cells (GCs), unipolar brush cells (UBCs), and glutamatergic CN neurons (Machold and Fishell, 2005; Wang et al., 2005; Englund et al., 2006; Yamada et al., 2014).

The transcriptomic data obtained in FANTOM5 reveal that in the cerebellum, *Pou3f1* expression peaks at E12, then gradually decreases as development progresses. As E12 is a key timepoint for the neurogenesis of several cerebellar cell types, *Pou3f1*’s temporal expression is notable and invites further investigation. Using *in situ* hybridization (ISH) and immunofluorescence (IF), we were able to determine to which lineage Pou3f1-expressing neurons belong, as well as the spatiotemporal characteristics of Pou3f1 expression in relation to those of other cell markers of the same lineage.

Our findings indicate that Pou3f1 is a marker for a previously unrecognized subtype of glutamatergic CN neurons that express neither Pax6 nor Tbr1. Importantly, our results illuminate the existence of other molecular programs instructing glutamatergic CN neuron development that are different from the “canonical” expression pathway of Atoh1→Pax6→Tbr1. Furthermore, glutamatergic CN neurons that arise from the RL are comprised of molecularly heterogeneous subpopulations, each likely exhibiting a distinct developmental regulatory program.

## Materials & Methods

### Mouse strains and husbandry

The mouse strain C57BL/6J was used for the characterization of wildtype (WT) Pou3f1 expression pattern during development.

The *Pax6* mutant strain, *Pax6*^*Sey*^ (originally obtained from Robert Grainger and Marilyn Fisher, University of Virginia), was bred as heterozygous pairs, phenotyped for eye sizes and presence of cataracts, and genotyped as previously described (Swanson et al., 2005). Experimental *Pax6*^*Sey*/*Sey*^ embryos were generated by intercrossing *Pax6*^*Sey/+*^ mice.

The *Atoh1-lacZ* reporter strain, *Atoh1*^*β-Gal*^ (obtained from Huda Zoghbi, Baylor College of Medicine), was genotyped by PCR according to the protocol previously described (Jensen et al., 2002). Experimental null embryos, *Atoh1*^*β-Gal/ β-Gal*^, were generated by intercrossing *Atoh1*^*β-*^ *Gal/+* mice.

All embryonic ages utilized in these experiments were confirmed based upon the appearance of a vaginal plug. The morning that a vaginal plug was detected was designated as E0.5. All studies were conducted according to the protocols approved by the Institutional Animal Care and Use Committee and the Canadian Council on Animal Care at the University of British Columbia

### Experimental mouse chimeras

Experimental mouse chimeras were generated as most recently described (Yeung et al., 2016), using the mating of heterozygous *Sey* mutants (Pax6^Sey/+^) to generate the mutant component of the chimeras. The WT component of the chimeras was generated from the mating of homozygous FVB.Cg-Tg(CAG-EGFP)B5Nagy/J mice (The Jackson Laboratory, Bar Harbor, ME; Stock number: 003516). Four-to-eight cell embryos from each of the mating schemes were brought into apposition in a small drop of culture medium and cultured overnight. On the next day, the successfully fused embryos were transferred to the uterine horn of pseudo-pregnant ICR females (plugged by a vasectomized male). All transplanted embryos were then collected at E18.5 and tail samples were processed for PCR to ascertain the Pax6 genotype as described previously (Swanson et al., 2005). Embryos were processed for histological analysis using a cryostat.

An estimate of percentage chimerism was made by the cells expressing GFP (the WT component of the chimera) in regions inside and outside of the cerebellum. For each chimeric brain, GFP expression from 13 to 16 coronal sections was analyzed and averaged.

CN neuron phenotype was assessed by counting Pou3f1^+^ cells from 13 to 16 coronal sections across the full cerebellum, right and left sides inclusive. We determined the number of Pou3f1^+^ CN neurons from the cerebella of two WT *Pax6*^*+/+*^ ↔ +/+ chimeras, three heterozygous *Pax6*^*Sey/+*^↔ +/+ chimeras, three mutant *Pax6*^*Sey/Sey*^ ↔ +/+ chimeras, and four mutant *Pax6*^*Sey/Sey*^ embryos. The total number of Pou3f1^+^ cells in each cerebellum was estimated, and averages were taken for all groups of embryos. For the mutant chimeric cerebellum, the expected number of Pou3f1^+^ cells was predicted based on the percent chimerism (of the WT and mutant genotypes) multiplied by the average cell counts from the WT and mutant cerebella, respectively (see text). Statistical significance between the expected and observed numbers of Pou3f1^+^ cells in the mutant chimeric cerebellum was determined by *chi*-square test.

### Tissue preparation and histology

Embryos were collected every day from E10.5 to E18.5. Neonatal pups were collected at P0 and P6. Embryos harvested between E10.5 to E15.5 were fixed by immersion in 4% paraformaldehyde in 0.1 M phosphate buffer (PB, pH 7.4) for 1 hour at 4°C. Embryos or pups harvested at E16.5 and later were transcardially perfused first with phosphate buffered saline (PBS), and then with 4% paraformaldehyde in 0.1 M PB. Brain tissues were isolated and further fixed in 4% paraformaldehyde in 0.1 M PB for 1 hour at room temperature. Fixed tissues were rinsed with PBS, followed by cryoprotection with 30% sucrose/PBS overnight at 4°C before embedding in Optimal Cutting Temperature (O.C.T.) compound (4583, Sakura Finetek USA). Tissues were sectioned in a cryostat at 12 μm thickness for IF and ISH, then cryosections were mounted on Superfrost™ slides (12-500-15, Fisher Scientific), air dried at room temperature, and stored at −80°C until used. Sagittal sections were cut from one side of the cerebellum to the other (left to right, or vice versa). Coronal sections were cut from the front of the cerebellum to the back. In all cases, observations were based on a minimum of 3 embryos per genotype per experiment.

### In situ hybridization

All ISH experiments were performed on tissues prepared as described above. Sense and antisense riboprobes corresponding to the cDNA fragment were synthesized and labeled with digoxygenin (DIG)-UTP. A cDNA library was obtained from E15.5 mouse brain using a cDNA synthesis kit (K1681, Thermo Scientific). Then the corresponding cDNA of *Pou3f1* was produced with this cDNA library, using the gene specific forward and reverse primers (Forward: 5’-AGCAGCGGAAGATCCAGAAT-3’; Reverse: 5’-TCGGTTTAGTCGGGCATACA-3’). These primers flank the 3’ untranslated region and yield a product of 1088bp; both the targeting region and length of the probe are important factors contributing to higher ISH specificity. Each of the resultant cDNA was cloned into the pGEM-T Easy vector (A1360, Promega) for the generation of cDNA templates. cDNA templates for the sense and antisense riboprobes were specifically made using the primers M13F: 5’-GTTTTCCCAGTCACGAC-3’ or M13R: 5’-CAGGAAACAGCTATGAC-3’ and the *Pou3f1*-specific forward or reverse primers. Riboprobes were produced using SP6 or T7 RNA polymerase (#EP0133 and #EP0111, Thermo Scientific, respectively) with the corresponding cDNA templates. The resultant riboprobes were precipitated using 5M ammonium acetate and 100% EtOH in RNase-free environment. Riboprobes were denatured at 72°C for 10 min, and incubated on ice for 5 min, then mixed with ULTRAhyb hybridization buffer (AM8670, Applied Biosystems) preheated at 68°C. Prior to hybridization, sections were acetylated with acetic anhydride in 0.1M triethanolamine at pH 8.0 and dehydrated with graded concentrations of RNase-free ethanol. Sections were first incubated with ULTRAhyb hybridization buffer at 68°C in a humid chamber for 1.5 hour, then replaced with riboprobe in ULTRAhyb hybridization buffer at 68°C overnight. After hybridization, the slides were rinsed with descending concentrations of salt: 4x SSC, 50% formamide in 2x SSC, and 2x SSC at 55°C. Sections were then treated with RNase A in 2x SSC at 37°C for 30 min, followed by 2x SSC, 1x SSC, and 0.5x SSC at 55°C. Afterwards, the sections were incubated with an anti-Dig antibody (11093274910, Roche) for 2 hours at room temperature. The slides were washed with maleic buffer, followed by reaction buffer, then the slides were colorized with NBT/BICP (11681451001, Roche). The reaction buffer was prepared according to manufacturer manual of NTP/BICP (11681451001, Roche). Following colorization, the slides were rinsed with 0.1M PB, then post-fixed in 4% paraformaldehyde, and washed with distilled water. The slides were dehydrated with graded concentrations of ethanol and xylene. At the end, cover slip was applied to the slide, using Paramount (SP15-500, Fisher Scientific) as the mounting medium.

### Immunofluorescence

For immunofluorescence, tissue sections were first rehydrated in PBS three times, five minutes each time, followed by a phosphate buffered saline with Triton X-100 (PBS-T) wash for five minutes. Sections were then incubated at room temperature for 1 hour with blocking solution (0.3% BSA, 10% normal goat serum, 0.02% sodium azide in PBS-T). Following the blocking step, the slides were subsequently incubated with primary antibody in incubation buffer (0.3% BSA, 5% normal goat serum, 0.02% sodium azide in PBS-T) at room temperature overnight in a humid chamber. Primary antibodies used were as follows: rabbit anti-Atoh1 (1:500; Proteintech, 21215-1 AP), rabbit anti-Brn2 (1:1000; Abcam, Ab94977), rabbit anti-Irx3 (1:8000; a gift from Tom Jessell, Columbia University), rabbit anti-Pax6 (1:200; Covance, PRB-278P, RRID:AB_291612), mouse anti-Pou3f1 (1:500; Millipore, MABN738), and rabbit anti-Tbr1 (1:800; Abcam, Ab31940, RRID:AB_2200219). Following the overnight incubation, the slides were rinsed in three PBS-T washes, 10 minute each time. The sections were then incubated with the corresponding secondary antibodies at room temperature for one hour, followed by three 0.1M PB washes and one 0.01M PB wash. Coverslips were applied to the slides using FluorSave mounting medium (345789, Calbiochem).

### Cell counts

We estimated the number of different classes of CN neurons by counting cells positive for the appropriate cell markers (Pou3f1, Brn2, or Irx3). Every 10th coronal section across the whole E18.5 cerebella was counted.

In all cases, observations were based on a minimum of 3 cerebella per genotype. Standardization of intensity, as well as cell counting, were performed using ImageJ’s built-in functions. Statistical significance between WT and *Pax6* mutant mice was determined by a One-Way ANOVA with post hoc test.

### Microscopy

Analysis and photomicroscopy were performed with a Zeiss Axiovert 200M microscope with the Axiocam/Axiovision hardware-software components (Carl Zeiss).

## Results

### Temporal and spatial expression patterns of Pou3f1 across developmental timepoints

The quantitative analysis of transcripts over time obtained through the 5’-CAGE analysis demonstrates that *Pou3f1* expression peaks at E12, then decreases as development proceeds (Fig. 1). To examine the spatial pattern of expression, we performed ISH and IF across multiple timepoints to complement the quantitative data. Our ISH and IF results illustrate that Pou3f1 is expressed prominently in cells along the SS between E10.5 and E12.5, but not in cells in the VZ (Fig. 2A, B, C). From E13.5 to E15.5, Pou3f1 is expressed in the nuclear transitory zone (NTZ), along with reduced Pou3f1 expression in the subpial stream (SS) (Fig. 2D). Between E18.5 and P6, Pou3f1 expression remains in the cerebellar white matter, with no noticeable change in expression pattern across time (Fig. 2E, F).

**Figure 1.**
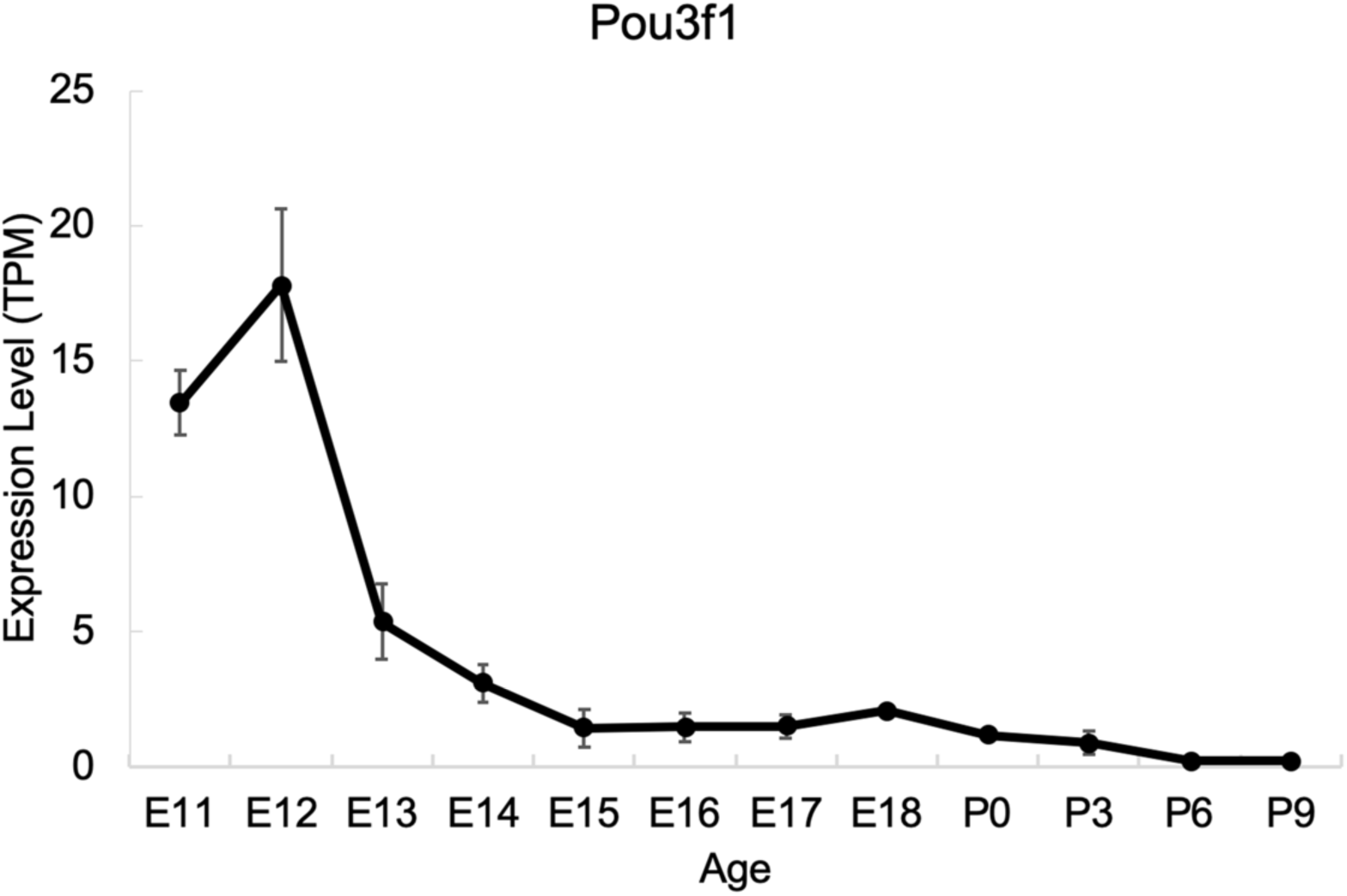
The transcriptional profile of *Pou3f1* during cerebellar development based upon the RIKEN FANTOM5 data. The *y*-axis shows expression level in transcripts per million (TPM). The *x*-axis represents developmental timepoints, from embryonic (E) to postnatal (P) ages. Error bars represent SE.

**Figure 2.**
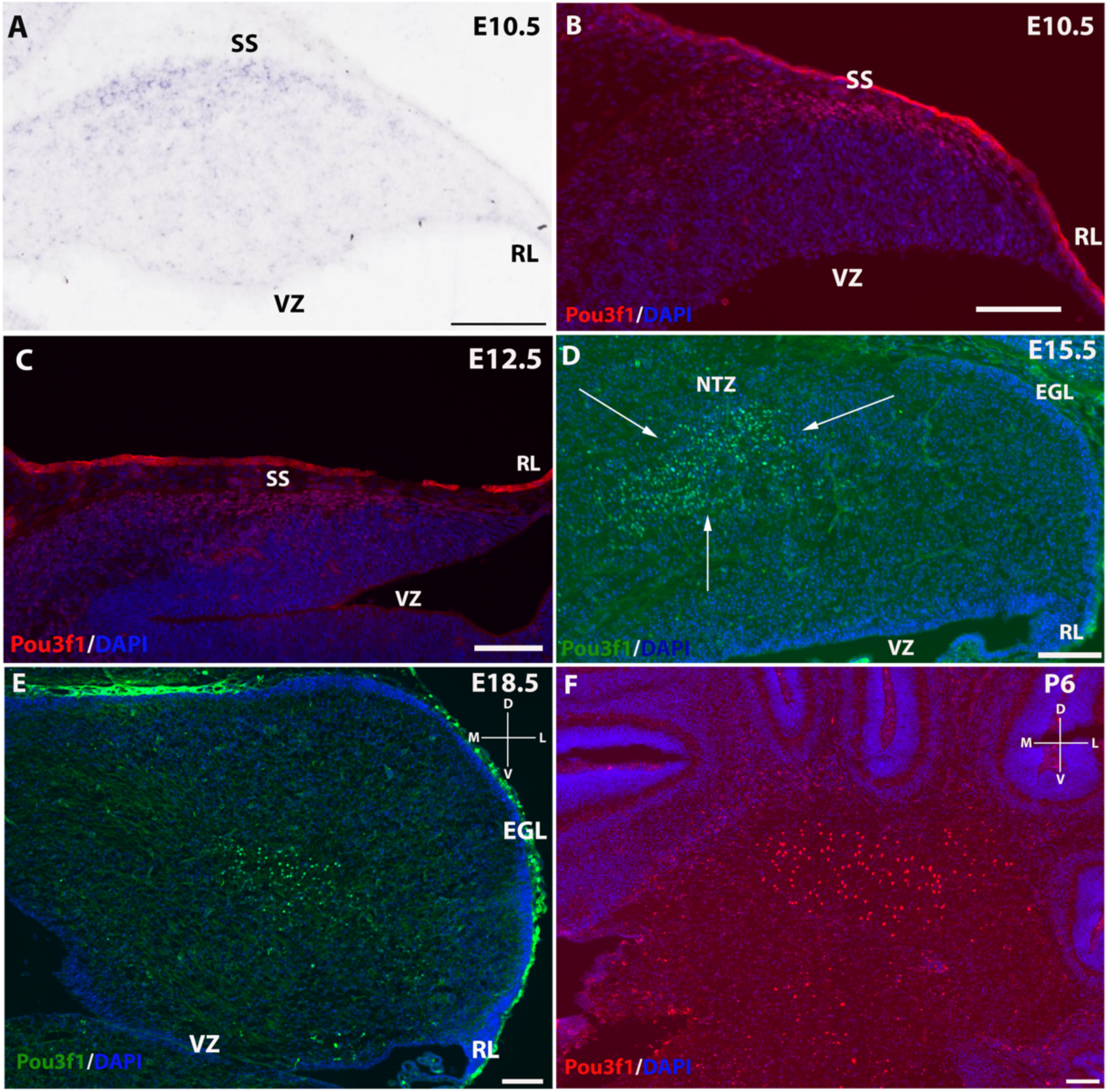
The expression pattern of Pou3f1 across space and time. **A, B, C**, In the E10.5 and E12.5 cerebella, *in situ* hybridization (ISH) (**A**) and immunofluorescent (IF) stainings (**B, C**) reveal that Pou3f1 is expressed in cells along the subpial stream (SS). **D**, In the E15.5 cerebellum, Pou3f1 is expressed primarily in the nuclear transitory zone (NTZ, white arrows). **E, F**, By E18.5, Pou3f1-expressing cells have reached the cerebellar white matter, with no noticeable change in spatial position at P6. **A** to **D** show sagittal sections of the cerebellum, while sections shown in **E** and **F** are in the coronal plane. D, dorsal. EGL, external germinal layer. L, lateral. M, medial. NTZ, nuclear transitory zone. RL, rhombic lip. SS, subpial stream. V, ventral. VZ, ventricular zone. Scale bars, 100 μm.

### Atoh1 and Pou3f1 demarcate three molecularly distinct regions in the developing cerebellum

The spatiotemporal expression pattern of Pou3f1 between E12.5 and P6 points to the possibility that Pou3f1^+^ cells are glutamatergic CN neurons that arise from the RL. To test this hypothesis, we examined the co-expression of Pou3f1 and Atoh1, the master transcription factor that is critical for the generation of glutamatergic populations of the cerebellum (Machold and Fishell, 2005; Wang et al., 2005).

At E12.5, IF co-staining of Pou3f1 and Atoh1 demonstrates that three molecularly distinct compartments are demarcated by the expressions of the two transcription factors (Fig. 3A): 1) Atoh1^+^ cells in the RL and the immediately adjacent segment of the SS, (Fig. 3B, C), 2) Pou3f1^+^cells in the main body of the SS (Fig. 3C, D), and 3) a transitional zone in the SS characterized by co-expression of Atoh1 and Pou3f1 (Fig. 3C). The three distinct compartments reveal an interesting region in which cells are both Pou3f1^+^ and Atoh1^+^. Moreover, the co-localization of Atoh1 and Pou3f1 confirms the hypothesis that Pou3f1^+^ cells are of glutamatergic lineage.

**Figure 3.**
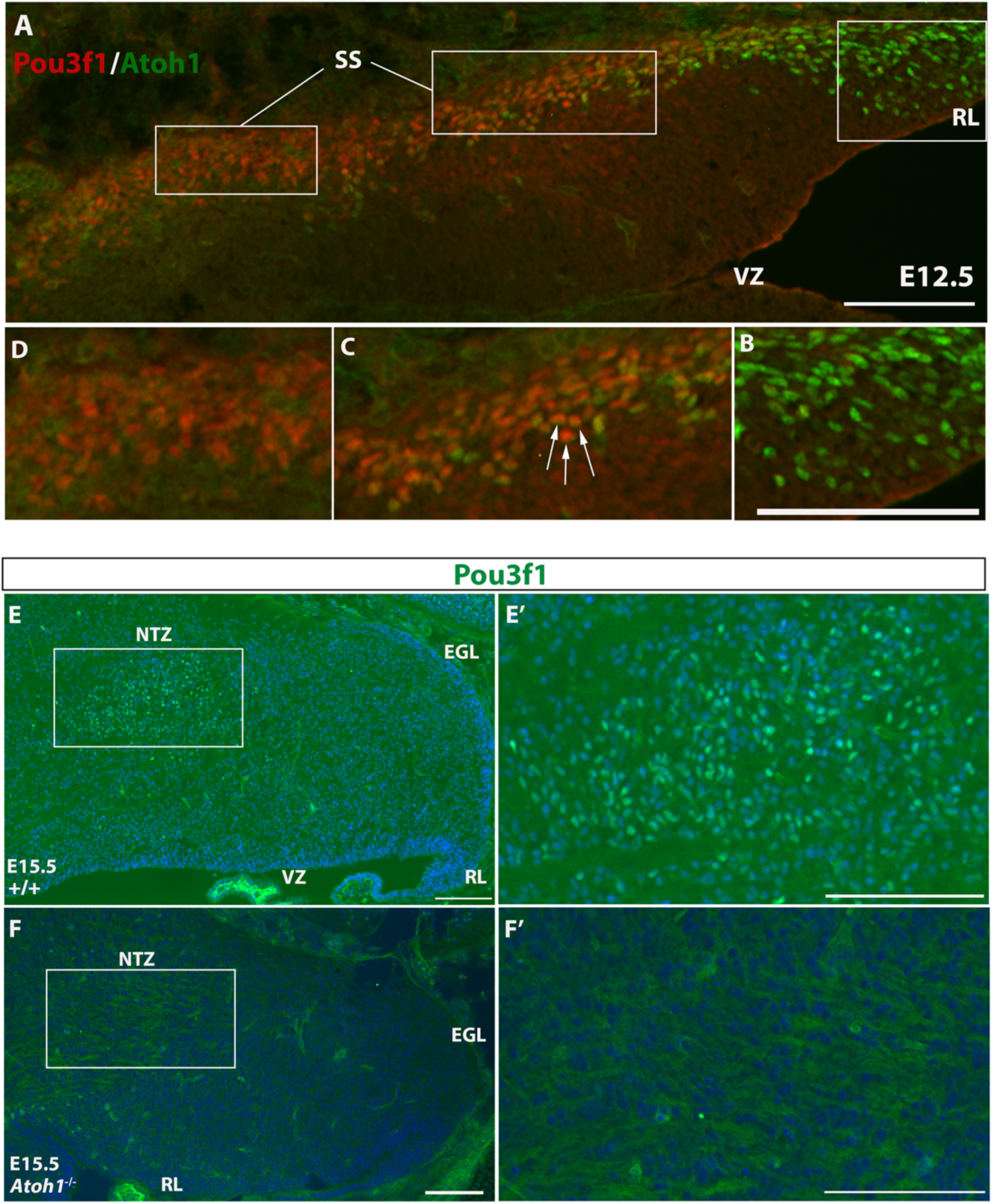
The expression patterns of Atoh1 and Pou3f1 in wildtype and *Atoh1*-null cerebella. **A**, Three molecular compartments are generated by Atoh1 and Pou3f1 expressions in the E12.5 cerebellar sagittal section. The rhombic lip (RL) is exclusively Atoh1^+^ (**B**), followed by a restricted region of the subpial stream (SS) characterized by co-localization of the two transcription factors (**C**, white arrows), and the rest of the SS marked exclusively by Pou3f1(**D**). **E, E’**, In the E15.5 wildtype cerebellar sagittal section, Pou3f1 expression is present in the nuclear transitory zone (NTZ). The highlight area in **E** is shown in **E’** in higher magnification. **F, F’**, However, in the corresponding *Atoh1*-null mutant cerebellar sagittal section, such expression is completely absent. Highlighted area in **F** is shown in **F’** in higher magnification. EGL, external germinal layer. NTZ, nuclear transitory zone. RL, rhombic lip. SS, subpial stream. VZ, ventricular zone. Scale bars, 100 μm.

### Pou3f1^+^ cells are absent from the Atoh1-null mutant cerebellum

Further evidence that Pou3f1 labels glutamatergic CN neurons comes from our examination of mice with homozygous knockout of *Atoh1*. In *Atoh1*-null mice, glutamatergic CN neurons are completely eliminated (Wang et al., 2005). Therefore, we expected Pou3f1^+^ neurons to be absent in the *Atoh1*-null cerebellum. In E15.5 WT cerebellar tissues, Pou3f1 is expressed in the NTZ (Fig. 3E, E’). However, in E15.5 *Atoh1*-null cerebellum, Pou3f1-expressing cells are undetectable, supporting the view that Pou3f1^+^ cells are Atoh1-dependent and in the glutamatergic lineage (Fig. 3F, F’). In short, investigation of the spatiotemporal relationships between Pou3f1 and Atoh1 suggests that Pou3f1^+^ neurons are glutamatergic. Moreover, the lack of Pou3f1 staining on *Atoh1*-null tissues provides evidence that Pou3f1 most likely acts downstream of Atoh1 in the context of glutamatergic CN neuron development.

### Pou3f1 and Pax6 label distinct cell populations

After validating that Pou3f1-expressing neurons are Atoh1-dependent and thus are glutamatergic, we investigated how other glutamatergic CN neuron markers might further define the molecular phenotype of Pou3f1-expressing cells. Pax6 is known to act downstream of Atoh1, and is important for the development of RL-derived cell types, including GCs, UBCs, and glutamatergic CN neurons (Fink et al., 2006; Yeung et al., 2016). Pax6 labels CN neurons along the SS, and as they enter the NTZ, Pax6 expression is downregulated (Fink et al., 2006; Yeung et al., 2016). Thus, the similarities in expression patterns between Pou3f1 and Pax6 are suggestive of co-expression. To test this possibility, we performed IF co-staining for Pou3f1 and Pax6 at E13.5 in WT cerebellar tissues, and surprisingly, the two genes are not co-expressed in the E13.5 cerebellum (Fig. 4A). Specifically, Pou3f1-expressing cells are localized to the NTZ (Fig. 4A’), while Pax6-expressing cells are restricted to the RL and the beginning of the SS (Fig. 4A’’). The absence of co-labeling between Pou3f1 and Pax6 at E13.5 caught our attention, as it was previously thought that the majority of glutamatergic CN neurons express Pax6 as part of their developmental progression (Yeung et al., 2016).

**Figure 4.**
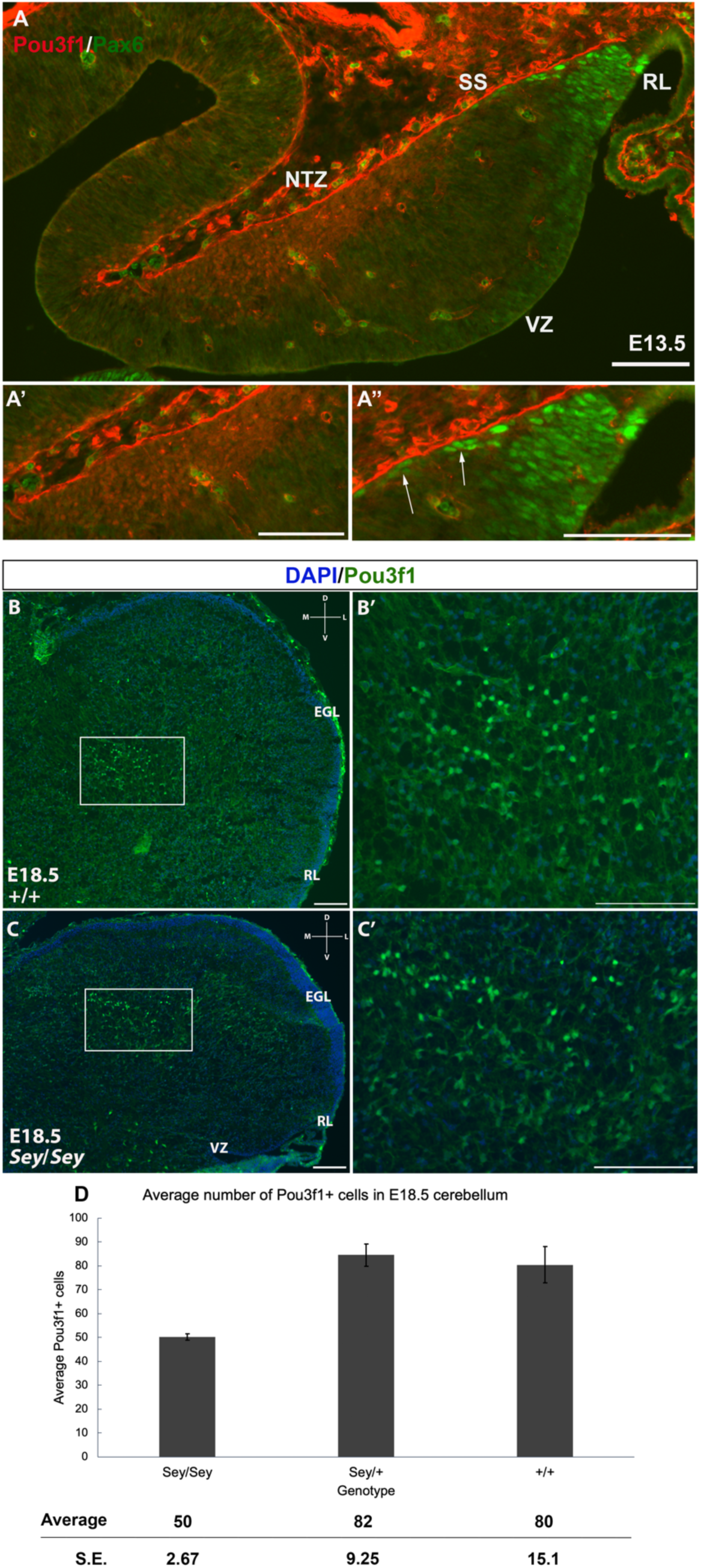
The absence of co-localization between Pou3f1 and Pax6 in wildtype cerebellum and the persistence of Pou3f1 expression in the Small Eye (*Sey*) mutant. **A**, In the E13.5 cerebellar sagittal section, lack of co-localization in expressions between Pou3f1 and Pax6 is evident, with Pou3f1-labeled cells predominantly in the nuclear transitory zone (NTZ) (**A’**), whereas Pax6-labeled cells are predominantly in the rhombic lip (RL) and the subpial stream (SS) (**A’’**, white arrows). **B, B’**, In the E18.5 wildtype cerebellar coronal section, Pou3f1-expressing cells are found to reside in the cerebellar white matter. The highlighted area in **B** is shown in **B’** in higher magnification. **C, C’**, In the E18.5 *Sey* cerebellar coronal section, Pou3f1 expression remains present. The highlighted area in **C** is shown in **C’** in higher magnification. **D**, One-Way ANOVA reveals that the average number of Pou3f1^+^ cells in the E18.5 *Sey* coronal section is significantly different from those in the heterozygous mutant and wildtype counterparts (*p* = 1.63 × 10^−2^). The average numbers were calculated by dividing the total number of Pou3f1^+^ cells in each animal by the total number of sides of cerebellar coronal sections counted. D, dorsal. EGL, external germinal layer. L, lateral. M, medial. NTZ, nuclear transitory zone. RL, rhombic lip. V, ventral. VZ, ventricular zone. Error bars represent SE. Scale bars, 100 μm.

### The persistence of Pou3f1 expression in the Sey mutant

To further explore the relationship between Pou3f1 and Pax6, we examined the expression of Pou3f1 in the *Pax6*-null *Se*y mutant. The *Sey* mutant has revealed that Pax6 plays a critical role in cerebellar development, as the *Sey* cerebellum suffers from improper foliation and abnormal morphology of the EGL (Engelkamp et al., 1999; Swanson et al., 2005); moreover, glutamatergic CN neurons and UBCs experience increased apoptosis (Engelkamp et al., 1999; Swanson et al., 2005). The lack of co-localization of Pou3f1 and Pax6 in the WT cerebellum suggests that these molecules operate independently and in separate cell populations. Thus, the *Sey* mutant serves as a good testbed to examine potential interactions between Pou3f1 and Pax6 in the development of glutamatergic CN neurons. As can be seen in Figure 4, Pou3f1 expression persists in the *Sey* mutant throughout development (Fig. 4B, B’, C, C’). To complement the qualitative observations of IF, we counted the number of Pou3f1^+^ cells in the cerebellum. Interestingly, at E18.5, One-Way ANOVA reveals that the average number of Pou3f1^+^ neurons in the *Sey* mutant cerebellum is significantly different from those in the heterozygous mutant and WT counterparts (*p* = 1.63 × 10^−2^) (Fig. 4D).

As previous studies indicated that Pax6 plays a non-cell autonomous role for the survival of glutamatergic CN neurons (Yeung et al., 2016), we tested the possibility of this effect of Pax6 on Pou3f1^+^ cells in experimental *Sey* chimeras (Fig. 5A). Experimental chimeras were composed of two genetically distinct cell lineages that were made by fusing two 4-8 cells blastocysts. Chimeric animals provide a unique opportunity to examine the impact of mutant cells in WT environment and vice versa. Three E18.5 *Sey* chimeras were obtained (see Methods), and were estimated to contain 0.5%, 10%, and 23% of WT *Pax6*-expressing cells from cerebellum, midbrain and brainstem tissues. We calculated the expected numbers of Pou3f1-labeled cells by multiplying the percentages of WT and *Pax6*^*-/-*^ cells by the average numbers of Pou3f1^+^ cells in the WT and *Sey*/*Sey* cerebella, respectively. The numbers of Pou3f1^+^ cells from the three chimeras were compared to their respective expected numbers using the *chi*-square test, to test the hypothesis that the number of Pou3f1^+^ neurons is in a linear relationship with the number of WT cells (Fig. 5B). The *chi*-square test rejects the null hypothesis (*p* = 4.83 × 10^−4^), thus indicating that there is not a linear relationship between the number of Pou3f1^+^ neurons and the percentage of WT cells in the cerebellum.

**Figure 5.**
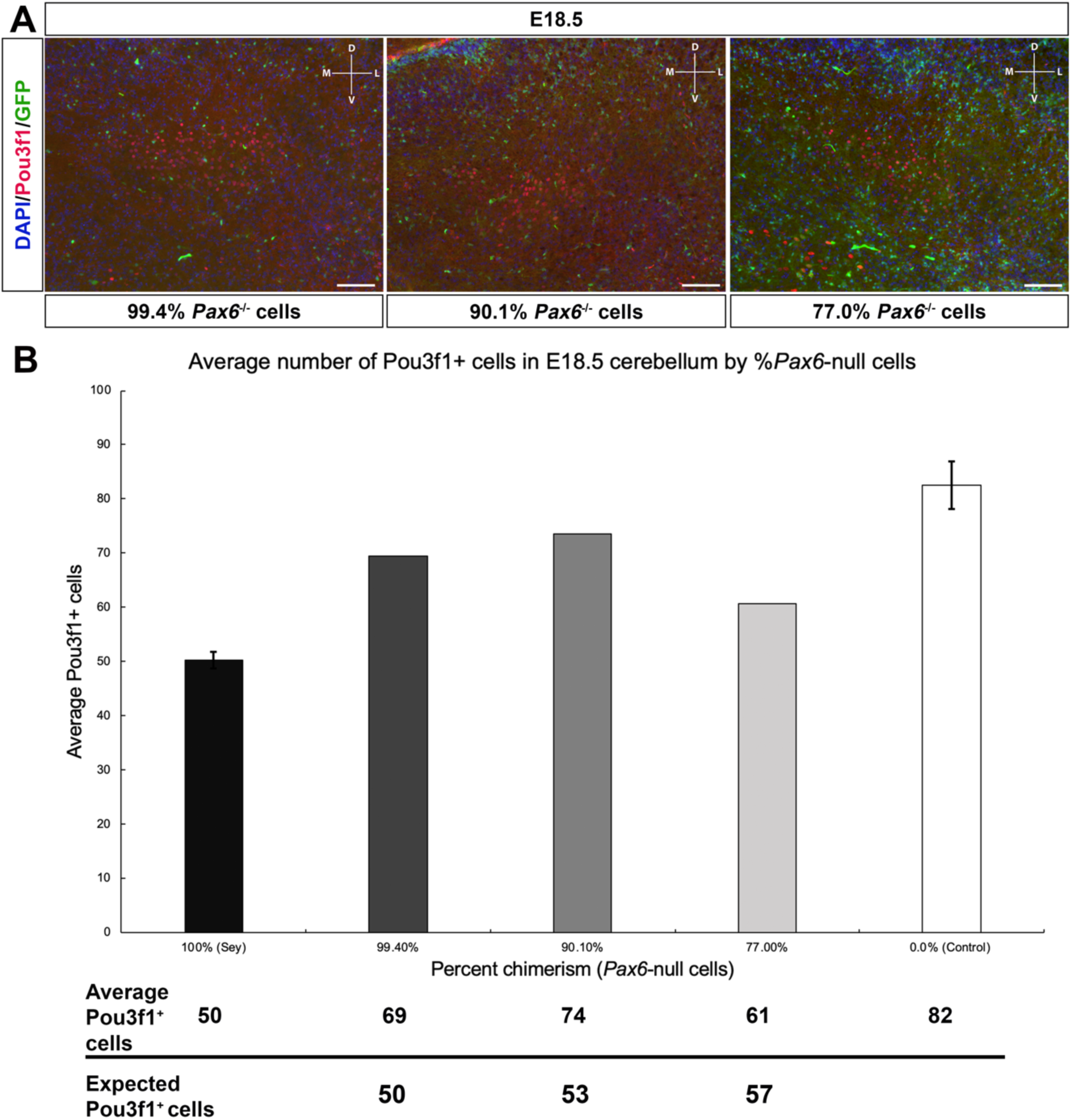
Pou3f1^+^ cells are present similarly in the Small Eye (*Sey*) mutant chimeras as in the *Sey*/*Sey* mutants. **A**, Pou3f1 expression in the E18.5 cerebellar coronal sections of *Sey* chimeras with different percentages of *Pax6*^-/-^ cells (decreasing percentages going from left to right). GFP labels wildtype cells. **B**, The average numbers of Pou3f1^+^ cells in the E18.5 cerebellar coronal sections were determined for each of the *Sey* chimeras, as well as for the *Sey/Sey* mutant and control counterparts (heterozygous and wildtype). The average numbers were calculated by dividing the total number of Pou3f1^+^ cells in each animal by the total number of sides of cerebellar coronal sections counted. The expected number of Pou3f1^+^ cells for each chimera animal was calculated by multiplying the corresponding percentages of wildtype and *Pax6*^-/-^ cells by the average numbers of Pou3f1^+^ cells in the wildtype and *Sey/Sey* cerebella, respectively. The *chi*-square test indicates that the observed numbers of Pou3f1^+^ cells in the *Sey* chimeras significantly deviate from the expected numbers (*p* = 4.83 × 10^−4^). D, dorsal. L, lateral. M, medial. V, ventral. Error bars represent SE. Scale bars, 100 μm.

### Pou3f1 defines a population of glutamatergic CN neurons distinct from the Tbr1^+^ population

Our study thus far has revealed that Pou3f1 labels CN neurons arising from Atoh1^+^ cells in the RL and diverging from the Pax6-expressing CN neurons. Next, we sought to determine how Pou3f1^+^ CN neurons map onto other known glutamatergic CN neuron populations, namely, those delineated by Tbr1 expression. Starting at E13.5, Tbr1 labels glutamatergic CN neurons in the NTZ that are destined to populate the fastigial nucleus, and its expression persists into adulthood in these neurons (Fink et al., 2006; Yeung et al., 2016). It is known that Tbr1 acts downstream of Atoh1 and Pax6, as Tbr1 is expressed in neither the *Atoh1*-null mice nor the *Sey* mutant (Fink et al., 2006; Yeung et al., 2016). To assess the relationship between Pou3f1^+^ and Tbr1^+^ CN neurons, we performed double labeling of Pou3f1 and Tbr1 across developmental timepoints. At all timepoints examined, Pou3f1 and Tbr1 are expressed by two non-overlapping populations. Namely, at E13.5, Pou3f1-expressing cells are localized more medially compared to those expressing Tbr1 (Fig. 6A). Then, at E15.5, even though Pou3f1^+^ CN neurons are found in the same medial-lateral plane as Tbr1^+^ CN neurons (Fig. 6B), it was observed that the majority of Pou3f1^+^ CN neurons are more rostral compared to those that are Tbr1^+^ (data not shown). Finally, at P6 (Fig. 6C), Pou3f1^+^ CN neurons populate the interposed and dentate nuclei (Fig. 6D and D’), lateral to the Tbr1^+^ counterparts that populate the fastigial nucleus (Fig. 6E, E’). The spatial distributions of Pou3f1^+^ and Tbr1^+^ populations in the P6 cerebellar coronal section are depicted in Fig. 6F.

**Figure 6.**
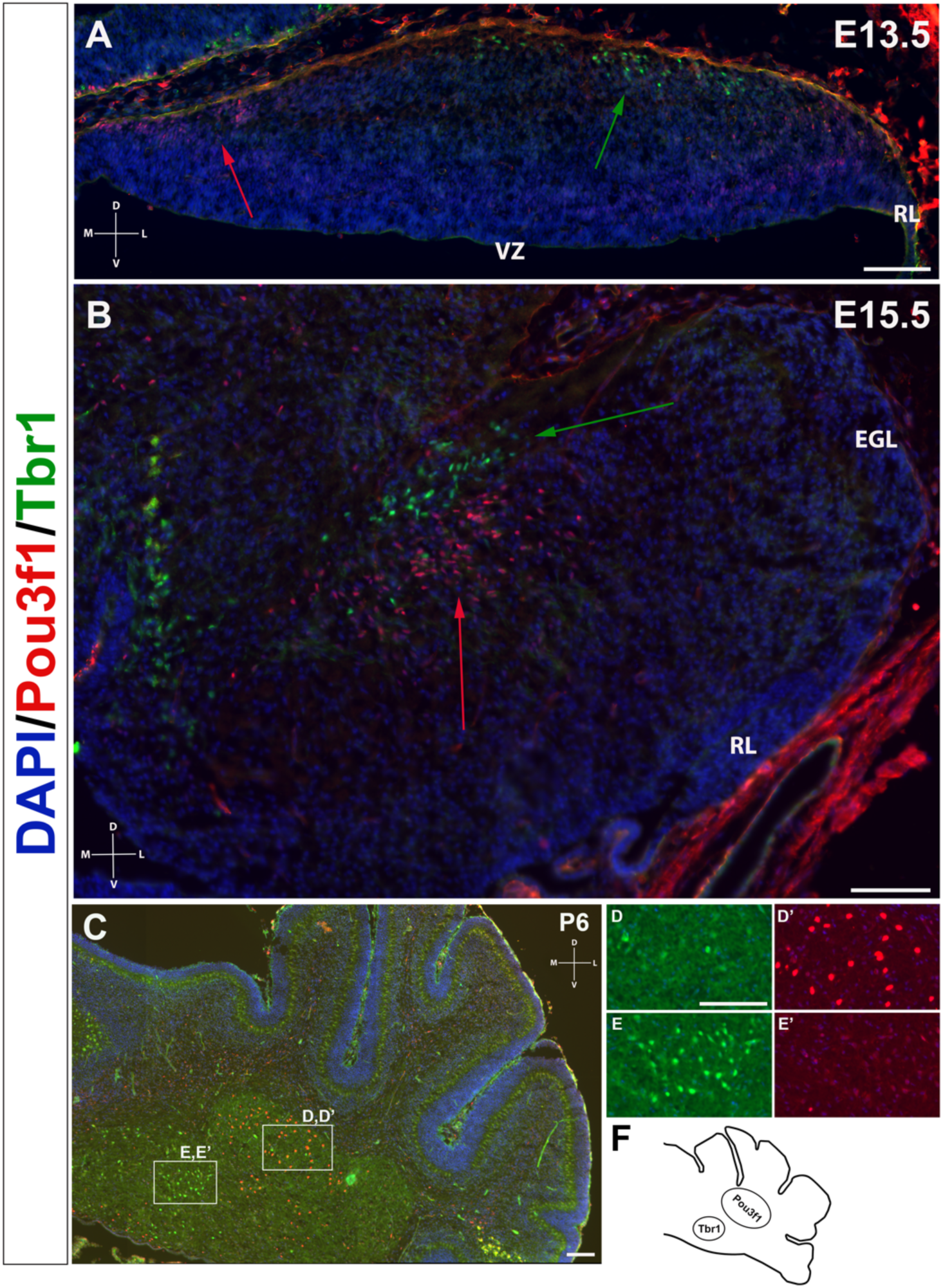
The non-overlapping expression patterns of Pou3f1 and Tbr1 throughout development. **A**, In the E13.5 wildtype cerebellar coronal section, Pou3f1-expressing cells are located more medially than Tbr1-expressing cells (red and green arrows, respectively). **B**, By E15.5, as shown in the wildtype coronal section, Pou3f1^+^ CN neurons and Tbr1^+^ CN neurons are found in the same plane along the medial-lateral axis (red and green arrows, respectively). **C**, At P6, the cluster of Pou3f1-labeled CN neurons is found more laterally than the Tbr1-labeled cluster, with the former residing primarily in the interposed nucleus (**D, D’**), while the latter residing in the fastigial nucleus (**E, E’**). The laterally highlighted area in **C** is shown in **D** and **D’** in higher magnification. Similarly, the medially highlighted area in **C** is shown in **E** and **E’** in higher magnification. **F**, A schematic diagram of the Pou3f1 and the Tbr1 clusters in the P6 cerebellar coronal section. D, dorsal. EGL, external germinal layer. L, lateral. M, medial. NTZ, nuclear transitory zone. RL, rhombic lip. V, ventral. VZ, ventricular zone. Scale bars, 100 μm.

The non-overlapping expression pattern between Pou3f1 and Tbr1 demonstrates that Pou3f1 defines a previously unidentified subtype of glutamatergic CN neurons that express neither Pax6 nor Tbr1 during their development. Moreover, by comparing the spatiotemporal pattern of Pou3f1 expression with that of Tbr1 expression, it is evident that as the cerebellum matures, Pou3f1^+^ CN neurons undergo a medial to lateral migration, while Tbr1^+^ CN neurons exhibit a lateral to medial migration during cerebellar development.

### Relationship of Pou3f1 with Brn2 and with Irx3

To further characterize the population of cells marked by Pou3f1, we examined its relationship with other markers of CN neurons. Brn2 has been shown to label CN neurons in the interposed and dentate nuclei (Fink et al., 2006). Co-staining of Pou3f1 and Brn2 on P6 coronal sections illustrates co-localization between these two molecules in both the interposed and dentate nuclei (Fig. 7A). The majority (approximately 70%) of Pou3f1^+^ CN neurons are also Brn2^+^ (Fig. 7B, B’, C, C’, white arrows). The P6 cerebellum also exhibits cells in the interposed and dentate nuclei that are singly labeled by either Pou3f1 (Fig. 7B, B’, C, C’, blue arrows) or Brn2 (Fig. 7B, B’, C, C’, orange arrows). As Brn2 expression is present in a larger proportion of interposed and dentate nuclei (Fig. 7A), these findings suggest that Pou3f1 molecularly defines a subtype within the Brn2-expressing CN neuron population. The spatial distributions of Pou3f1^+^, Tbr1^+^, and Brn2^+^ populations in the P6 cerebellar coronal section are depicted in Fig. 7D.

**Figure 7.**
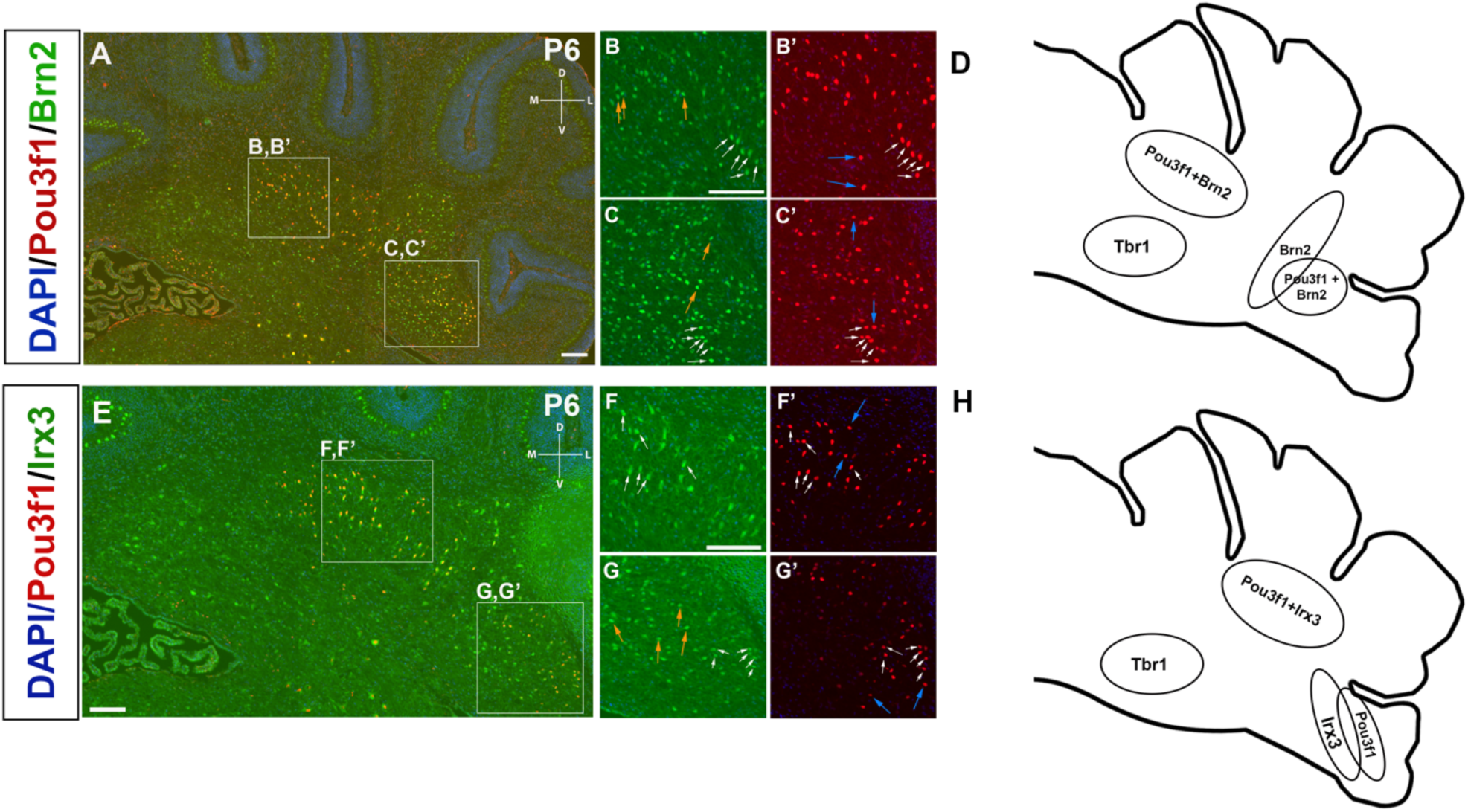
Characterization of Pou3f1 expression with Brn2 expression and with Irx3 expression in the P6 cerebellar coronal sections. **A**, Co-expression of Pou3f1 and Brn2 is evident in both the interposed (**B, B’**) and the dentate nuclei (**C, C’**). Approximately 70% of Pou3f1^+^ CN neurons also express Brn2 (white arrows). Blue arrows indicate cells that are Pou3f1^+^ but Brn2^-^, while orange arrows indicate cells that are Brn2^+^ but Pou3f1^-^. **D**, A schematic diagram showing the spatial distributions of Pou3f1^+^, Tbr1^+^, and Brn2^+^ populations in the P6 cerebellar coronal section. **E**, Co-expression of Pou3f1 and Irx3 is evident predominantly in the interposed nucleus (**F, F’**), with minor co-expression in the dentate nucleus (**G, G’**). Approximately 55% of Pou3f1^+^ CN neurons also express Irx3 (white arrows). Blue arrows indicate cells that are Pou3f1^+^ but Irx3^-^, while orange arrows indicate cells that are Irx3^+^ but Pou3f1^-^. **H**, A schematic diagram showing the spatial distributions of Pou3f1^+^, Tbr1^+^, and Irx3^+^ populations in the P6 cerebellar coronal section. D, dorsal. L, lateral. M, medial. V, ventral. Scale bars, 100 μm.

Another cell marker, Irx3, has been shown to label CN neurons emerging from both the VZ and the RL (Morales and Hatten, 2006). We found that in the P6 cerebellum (Fig. 7E), more than half (∼ 55%) of Pou3f1^+^ CN neurons also express Irx3 (Fig. 7F, F’, G, G’, white arrows). Besides the Pou3f1^+^/Irx3^+^ population, CN neurons singly labeled by either Pou3f1 (Fig. 7F, F’, G, G’, blue arrows) or Irx3 (Fig. 7F, F’, G, G’, orange arrows) are also present. Analogous to the results with Brn2, Irx3 is expressed in a larger proportion of CN neurons (Fig. 7E). These findings suggest that Pou3f1 expression defines a molecular subpopulation within the Irx3-labeled cohort. The spatial distributions of Pou3f1^+^, Tbr1^+^, and Irx3^+^ populations in the P6 cerebellar coronal section are depicted in Fig. 7H.

As might be expected from the presence of Pou3f1 cells in the *Sey* mutant, Brn2 and Irx3 remain robustly expressed in the E18.5 *Sey* mutant cerebellum. In particular, similar to the expression patterns in WT cerebellum (Fig. 8A, B), both Brn2 and Irx3 label CN neurons in the interposed and dentate nuclei (Fig. 8A’, B’).

**Figure 8.**
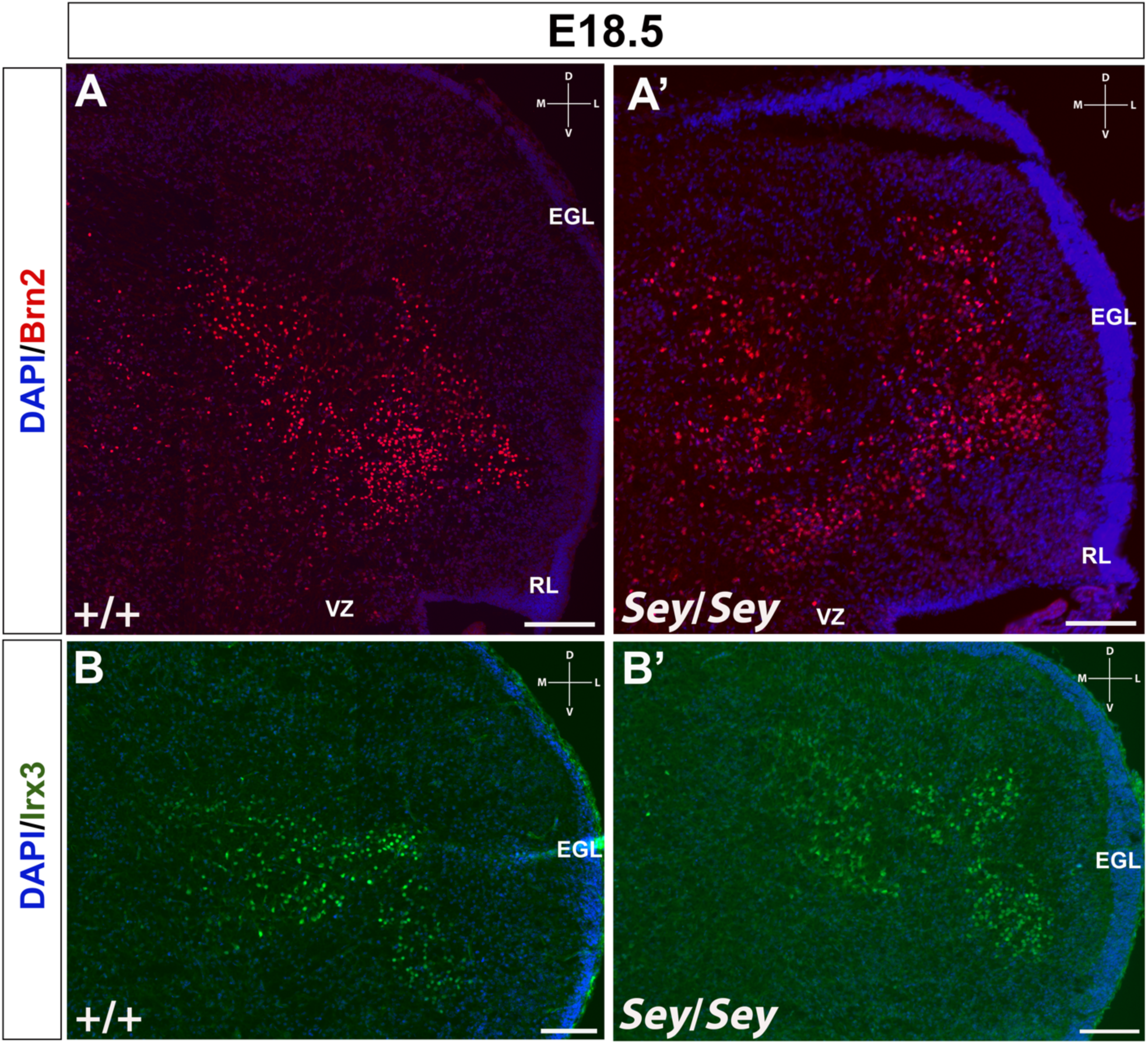
Brn2 and Irx3 remain expressed in the Small Eye (*Sey*) mutant. **A**, In the E18.5 wildtype cerebellar coronal section, Brn2 is robustly expressed in both the interposed and dentate nuclei. **A’**, Brn2 expression in the E18.5 *Sey* mutant cerebellar coronal section recapitulates that in the wildtype counterpart. **B**, Similarly, in the E18.5 wildtype cerebellar coronal section, Irx3 is expressed in both the interposed and dentate nuclei. **B’**, Irx3 expression in the E18.5 *Sey* mutant cerebellar coronal section recapitulates that in the wildtype counterpart. D, dorsal. EGL, external germinal layer. L, lateral. M, medial. RL, rhombic lip. V, ventral. VZ, ventricular zone. Scale bars, 100 μm.

## Discussion

The present study has demonstrated that Pou3f1 is a marker for a previously unrecognized subtype of glutamatergic CN neurons that diverge from the Pax6-lineage during cerebellar development. This exciting finding sheds light on the view that the glutamatergic CN neuron population is composed of at least two molecularly heterogeneous subpopulations, specified by distinct molecular programs.

### A molecular hand-off between Pou3f1 and Atoh1 during glutamatergic CN neuron development

Our examination of Pou3f1 and Atoh1 expressions in E12.5 cerebellum delineates a molecular hand-off between Pou3f1 and Atoh1 as cells migrate out of the RL. This process of a hand-off is evident in the portion of the SS directly adjacent to the RL where the two transcription factors co-localize. The downregulation of Atoh1 and the concomitant upregulation of Pou3f1 happening during this hand-off define the development of CN progenitor cells that emerge from the RL. Specifically, the degree of Pou3f1 expression indicates the developmental state of glutamatergic CN neurons: 1) the newly generated CN neuron progenitors are Atoh1^+^ only, 2) followed by cells that co-express Pou3f1 and Atoh1, and 3) the most mature glutamatergic CN neurons are exclusively Pou3f1^+^.

Mechanistically, the molecular hand-off between Pou3f1 and Atoh1 may be mediated at the level of transcriptional regulation. Namely, analogous to the regulatory model that governs mouse motor neuron differentiation proposed by Rhee et al. (2016), we suspect that, as cells leave the RL, the enhancer driving Atoh1 expression at the RL is transcriptionally silenced. This silencing event frees up a yet to be identified transcription factor, now able to bind to another enhancer that drives the expression of Pou3f1 along the SS. This *cis* regulatory change may work in conjunction with a *trans* regulatory change, involving a switch in binding partner with the common transcription factor governing both Pou3f1 and Atoh1 expressions, such that the entire transcriptional complex exhibits varying affinities for different enhancers in a stage-specific manner during glutamatergic CN neuron development. A suggestion of a similar event has been previously described in the E15.5 RL between Wls^+^ and Atoh1^+^ cells (Yeung et al., 2014).

### Uncovering a newly defined program to become glutamatergic CN neurons

The current view of glutamatergic CN neuron development entails the sequential expression of transcription factors Atoh1→Pax6→Tbr1, based upon gene expression and knockout studies (Wang et al., 2005; Fink et al., 2006; Yeung et al., 2016). However, our present findings have illustrated that besides this “canonical” pathway, there exists other molecular programs involved in CN neuron development, whose molecular hallmarks diverge after Atoh1 expression.

The classification of Pou3f1-expressing CN neurons as a distinct subclass that is independent of the canonical pathway, has motivated us to determine other molecular signatures of Pou3f1^+^ cells, in hopes of identifying a new expression program that governs the development of Pou3f1-expressing CN neurons. In the present work, we examined the expressions of Brn2 and Irx3, whose lineage associations still need to be determined. Brn2 is shown to label both interposed and dentate CN neurons (Fink et al., 2006). As described in our results, it is evident that cells can be positive for either Brn2 or Pou3f1 alone, which indicates that Brn2 and Pou3f1 can be expressed independently of each other. Moreover, because Brn2 is expressed in a greater proportion of CN neurons in the interposed and dentate nuclei compared to Pou3f1 expression, while our present work has characterized Pou3f1 as an exclusive marker of glutamatergic CN neurons, it is possible that within the entire population of Brn2^+^ cells, those that are singly labeled by Brn2 are GABAergic, while those co-expressing Pou3f1 are glutamatergic. Irx3 is another CN neuron marker documented to demarcate cells that migrate from both the SS and the VZ (Morales and Hatten, 2006). Analogously, within the population of Irx3^+^ CN neurons, cells singly labeled by Irx3 are likely GABAergic, while those co-expressing Pou3f1 are likely glutamatergic.

By comparing the glutamatergic component of Brn2^+^ population with that of the Irx3^+^ population, two possible models for the number of subtypes of Pou3f1-expressing CN neurons emerge (Figure 9): 1) Pou3f1 marks *two* subtypes of glutamatergic CN neurons; in particular, cells that are Pou3f1^+^/ Brn2^+^/Irx3^+^, and cells that are Pou3f1^+^ only (Fig. 9A); 2) Pou3f1 marks *three* subtypes of glutamatergic CN neurons; namely, cells that are Pou3f1^+^/Brn2^+^/Irx3^-^, cells that are Pou3f1^+^/Brn2^-^/Irx3^+^, and cells that are Pou3f1^+^ only (Fig. 9B). Answering this question will potentially illuminate a new paradigm that different subpopulations of CN neurons are delineated by the various combinatorial expressions of Pou3f1, Brn2, and Irx3.

**Figure 9.**
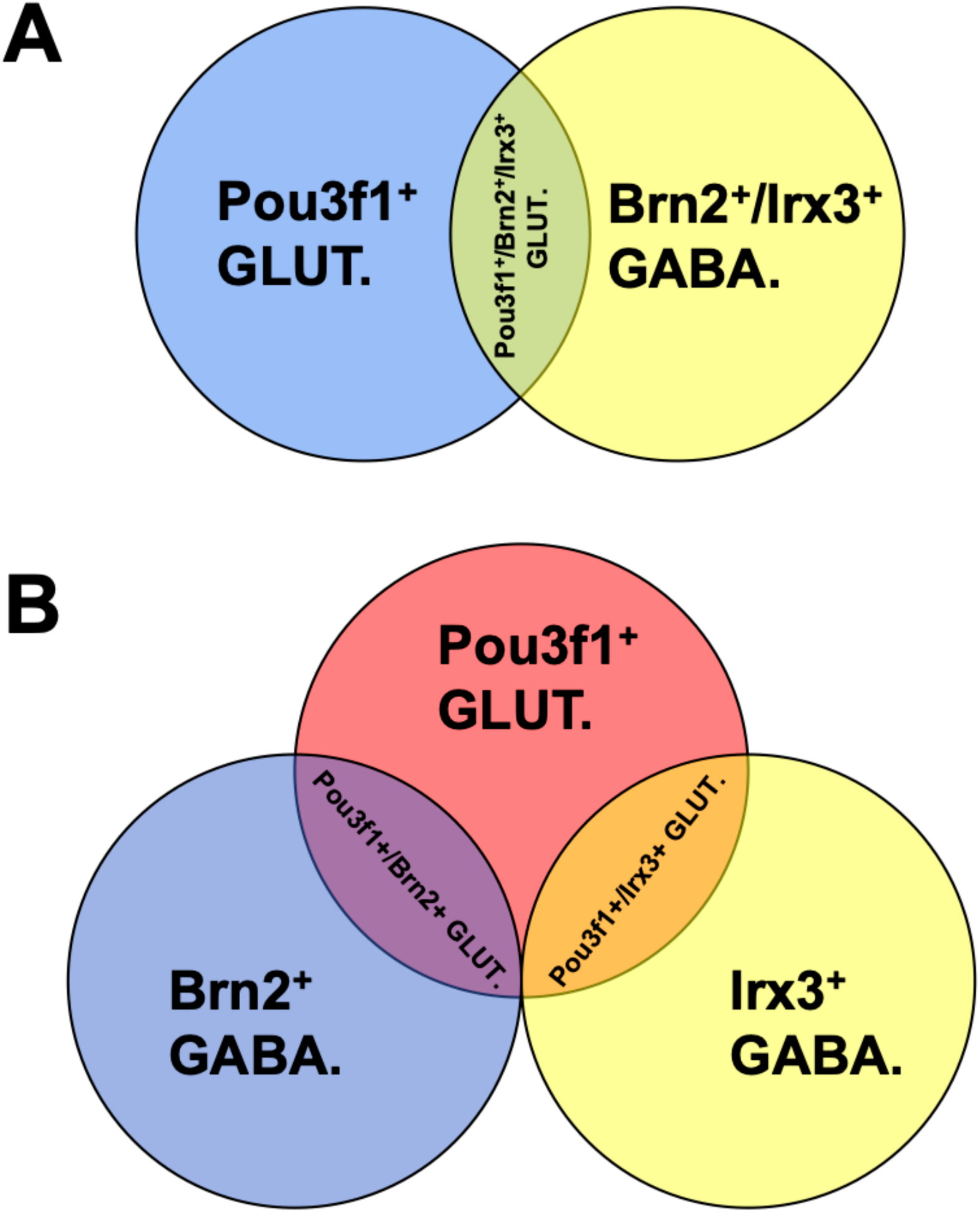
Two possible models for the number of subpopulations of glutamatergic CN neurons marked by Pou3f1. **A**, If the glutamatergic subpopulation delineated by Brn2 is the same as that delineated by Irx3, Pou3f1 marks at least *two* molecularly distinct subtypes of CN neurons: cells that are Pou3f1^+^/Brn2^+^/Irx3^+^, and cells that are Pou3f1^+^ only. **B**, On the other hand, if the two glutamatergic subpopulations (i.e. Brn2^+^ and Irx3^+^) are *not* the same, then Pou3f1 marks at least *three* molecularly distinct subtypes of CN neurons: cells that are Pou3f1^+^/Brn2^+^/Irx3^-^, cells that are Pou3f1^+^/Irx3^+^/Brn2^-^, and cells that are Pou3f1^+^ only. *Note: the areas of overlap between the circles are not proportional to the proportions of CN neurons that are labeled by the indicated markers*.

The observation that both Brn2 and Irx3 expressions persist in the E18.5 *Sey* mutant cerebellum suggests that Brn2 and Irx3 expressions are also independent of the canonical cascade. This key finding leads to a novel hypothesis that Pou3f1, Brn2, and Irx3 may all be part of a novel “non-canonical” molecular program involved in the development of a previously unidentified subtype of glutamatergic CN neurons, distinct from the canonical molecular cascade of Atoh1→Pax6→Tbr1 (Fig. 10).

**Figure 10.**
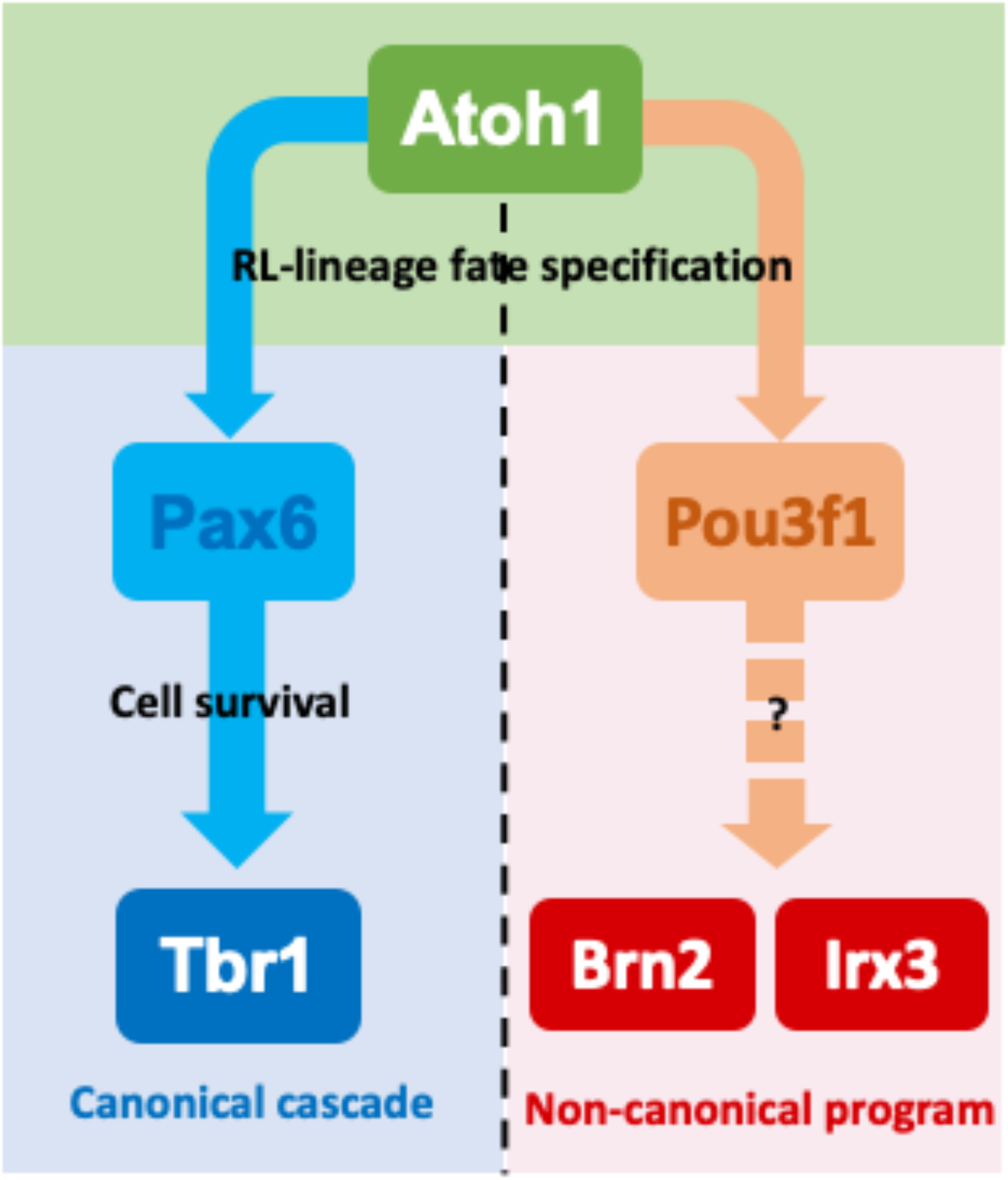
Two molecular models of glutamatergic CN neuron development. Atoh1 is expressed in all glutamatergic CN neuron progenitors arising out of the rhombic lip (RL) at E10.5. Then, for the canonical cascade, beginning in E13.5, the progenitors along the subpial stream (SS) begin to express Pax6. As these cells enter the nuclear transitory zone (NTZ), Pax6 expression is downregulated, while Tbr1 expression is upregulated. In the proposed non-canonical molecular program, Pou3f1 marks the first step of divergence from the canonical pathway, followed by Brn2 and Irx3 expressions. There may be additional molecules that are turned on between the onset of Pou3f1 expression and the onsets of Brn2 and Irx3 expressions. Unlocking this novel molecular program will help reveal the potential existence of other molecularly distinct subtypes of glutamatergic CN neurons.

However, it is notable that despite the divergence of Pou3f1^+^ CN neurons from the canonical Pax6-dependent pathway, we did observe a significant reduction of the number of Pou3f1^+^ cells in the *Sey* mutant cerebellum. This result suggests that a role, to be defined, exists for Pax6 in influencing the number of Pou3f1-expressing CN neurons. This possibility of the influence of Pax6 is also supported in our *Sey* chimera study, in which the numbers of Pou3f1^+^ cells are reduced in a non-linear fashion with increasing percentage of *Pax6*^*-/-*^ chimerism, thus pointing to an indirect role of Pax6 in regulating the number of another subtype of glutamatergic CN neurons. Elucidating this indirect modulatory relationship between Pou3f1 and Pax6 will undoubtedly further our understanding of glutamatergic CN neuron development.

### Pre-existing datasets provide insight into the molecular blueprint for CN neuron development

Previous ChIP-seq analysis of Pou3f1 performed in E4 mouse ESCs found that both *Brn2* and *Irx3* are transcriptional targets of Pou3f1 (Zhu et al. 2014). Moreover, when *Pou3f1* is overexpressed, both *Brn2* and *Irx3* are significantly upregulated (*p* = 7.91 × 10^−3^ and *p* = 7.04 × 10^−15^, respectively). These findings suggest that Pou3f1 most likely acts upstream of and positively regulates the expressions of Brn2 and Irx3. These results correlate with our observation that Pou3f1 is expressed earlier than Irx3 in glutamatergic CN neurons. Namely, Pou3f1 expression is detected as early as E10.5 along the SS, while Irx3 expression is detected not until E12.5 (data not shown).

Single-cell RNA sequencing (scRNA-seq) is a valuable technique to determine the expression profiles of specific genes at single-cell resolution. Recently published scRNA-seq data found that Pou3f1 expression is associated with postmitotic NTZ neurons, in which Atoh1 expression is also detected (Vladoiu et al. 2019). This result is consistent with our characterization of Pou3f1’s expression pattern. In addition, scRNA-seq also demonstrates that Brn2 and Irx3 expressions are found in both glutamatergic and GABAergic cell types (Vladoiu et al., 2019). This is aligned with the idea that within the Brn2^+^ and Irx3^+^ populations, those that are Pou3f1^-^ are GABAergic, while those co-expressing Pou3f1 are glutamatergic.

### Correlating the molecular heterogeneity of the cerebellar nuclear progenitor populations with their eventual functions

The CN neuron population consists of a heterogeneous population of cells: the excitatory glutamatergic CN neurons, as well as the inhibitory GABAergic and glycinergic CN neurons (Chen and Hillman, 1993). Glutamatergic CN neurons can be further divided into subgroups based on their mediolateral placement within the cerebellar white matter. While it is clear that excitatory and inhibitory CN neurons arise from RL and VZ, respectively, how do the cells that arise from the same germinal zones acquire different subtype identities is unknown. Are the glutamatergic CN neuron progenitors all generated following the same program then undergo subsequent specifications later in neuronal maturation? Or are cell fate specifications early events? Our data support the latter possibility, in which once glutamatergic CN neuron progenitors depart from the RL, the Pou3f1-program is activated and it identifies cells of the interposed and dentate nuclei. In contrast, the Tbr1^+^ fastigial CN population is generated following the canonical Atoh1→Pax6→Tbr1 pathway. Given our current understanding of the anatomical compartmentalization of CN neurons and their projections to other brain regions, the distinct molecular programs (Pax6-vs. Pou3f1-programs) come with functional differences. The fastigial CN (Tbr1^+^) neurons project to vestibular and reticular nuclei and are involved in control of head and ocular movements (Ruigrok et al, 2015). On the other hand, the interposed CN neurons (Pou3f1^+^/Irx3^+^/Brn2^+^) project to the red nucleus, which controls sensory and motor processing. The dentate CN neurons (Pou3f1^+^/Irx3^+^/Brn2^+^) project to the thalamus and cerebral cortex (Ruigrok et al, 2015), which play key roles in complex sensorimotor coordination and executive functions. Within the interposed or dentate nuclei, our present work reveals subpopulations characterized by Pou3f1 expression that are further defined by the combination of either Irx3 or Brn2 expressions. As Purkinje cells that innervate the CN neurons are also compartmentalized biochemically (e.g. Aldolase C/Zerbin II) (Brochu et al, 1990), it will be interesting to know whether there is a correlation in molecularly defined subtype identities between CN neurons and Purkinje cells.

## Acknowledgments

This work was supported by the Natural Sciences and Engineering Research Council (NSERC) of Canada and the Howard Hughes medical institute (HHMI). We thank Remi Robert for the assistance in bioinformatic analysis. We also thank Miguel Ramirez, Ishita Gupta and Eric Chow for their technical assistance.

